# Unsupervised Detection of Rare Events in Liquid Biopsy Assays

**DOI:** 10.1101/2025.01.29.635501

**Authors:** Javier Murgoitio-Esandi, Dean Tessone, Amin Naghdloo, Stephanie N. Shishido, Brian Zhang, Haofeng Xu, Agnimitra Dasgupta, Jeremy Mason, Rajiv M. Nagaraju, James Hicks, Peter Kuhn, Assad Oberai

## Abstract

The use of liquid biopsies in the detection, diagnosis and treatment monitoring of different types of cancers and other diseases often requires identifying and enumerating instances of analytes that are rare. Most current techniques that aim to computationally isolate these rare instances or events first learn the signature of the event, and then scan the appropriate biological assay for this signature. While such techniques have proven to be very useful, they are limited because they must first establish what signature to look for, and only then identify events that are consistent with this signature. In contrast to this, in this study, we present an automated approach that does not require the knowledge of the signature of the rare event. It works by breaking the assay into a sequence of components, learning the probability distribution of these components, and then isolating those that are rare. This is done with the help of deep generative algorithms in an unsupervised manner, meaning without a-priori knowledge of the rare event associated with an analyte. In this study, this approach is applied to immunofluorescence microscopy images of peripheral blood, where it is shown that it successfully isolates biologically relevant events in blood from normal donors spiked with cancer-related cells and in blood from patients with late-stage breast cancer.

## 1. Introduction

Liquid biopsy (LBx) has demonstrated the feasibility and clinical utility of blood-based cancer detection through applications in early detection, disease monitoring, and treatment management (1–7). Studies have shown that even asymptomatic patients can exhibit detectable levels of cancer-associated analytes in the blood (8–12). These analytes include acellular components such as cell-free DNA, RNA, proteins, extracellular vesicles, and cellular components like circulating cancer cells and tumor microenvironment cells. While cell-based detection approaches have been shown to identify a wide spectrum of cancer-related cells, they may struggle to scale into clinical practice due to the high degree of human involvement required for evaluating each assay in order to identify these rare events.

Circulating tumor cell (CTC) counts have been demonstrated to have prognostic value (1–3) and predictive utility (4, 13–15), while CTC characterization has shown substantial heterogeneity in both phenotype (16–19) and genotype (20–23).

Specific biological features, such as protein marker expression, have been found to be critical for therapeutic decisionmaking(4, 7). However, the field has been limited to either enumeration approaches of CTCs in clinical trials or limited biological characterization in clinical studies. While enumeration approaches have demonstrated clinical utility, biological characterization connects primary tumors to metastatic disease in ways that could offer deeper clinical insights.

Several sample preparation methods have been developed (1, 2, 24), each of which produces image datasets of target cells (cancer-related cells) mixed with non-target immune cells, often at ratios as extreme as 1 in 1 million. These imaging results require extensive human interpretation, typically performed by a pathology-trained technician supported by computational algorithms, which require significant prior knowledge about features that are biologically relevant. This restricts the scalability across multiple disease systems and laboratories.

Beyond scalability limitations, the heterogeneity of biomarkers emerging from LBx highlights the need for more generalizable analyte classification and discovery tools. Within the cancer cell population, various phenotypes—including platelet-coated CTCs (7) (CTCs that have platelets attached), epithelial-to-mesenchymal transition (EMT) CTCs (25) (cells transitioning from an epithelial to a mesenchymal state), and CTC clusters (26–30) (aggregates of CTCs)—have emerged as powerful predictive biomarkers in prostate, breast, lung, colorectal, and other cancers. Additionally, increasing evidence has demonstrated the presence of various tumor microenvironment cells in the blood of cancer patients at clinically relevant levels, including circulating endothelial cells (31) and cancer-associated fibroblasts (32), which can serve as companion biomarkers to traditional CTCs. Methods that enrich for a specific cellular population limit the ability to detect the heterogeneity of known circulating cancer-associated cells and to discover novel biomarkers in the LBx. Further, if multiple classes of events are deemed important, methods that can detect each class must be developed, which can be a difficult task as it requires large amounts of labeled data. These factors necessitate approaches that can accommodate biomarker diversity without relying on significant prior knowledge. With this as motivation, we present an automated, unsupervised approach that does not require the prior specification or knowledge of a relevant or interesting event. Instead, the approach operates under the principle that these events tend to be rare, and then develops a method for identifying a small cohort of the most rare events without any supervision regarding what these rare events are.

In machine learning, the task of identifying rare events is often referred to as anomaly detection. Unsupervised anomaly detection is carried out without any prior knowledge regarding which events are rare and is accomplished by two broad categories of techniques. The first includes methods that explicitly evaluate the probability density (or log-density) of a given sample. This is done by transforming the sample of interest from its native probability measure to a known, reference measure, and computing the Jacobian of this transformation. The transformation may be achieved by energybased models (EBMs) (33), normalizing flows (NFs) (34), and score-based diffusion models (35). For an application of these models to anomaly detection the reader is referred to (36, 37). The evaluation of the probability (or log-probability) typically requires computing the Jacobian of the transformation, which makes these techniques computationally expensive.

The second category of anomaly detection methods includes those that train an autoencoder (AE) to reproduce events from the distribution of interest, and then use the reconstruction error as a metric of rarity (38–40). AEs are a class of generative, unsupervised learning models with two components: an encoder and a decoder. The encoder network reduces the dimensionality of the input data to an n-dimensional vector (latent vector), and the decoder network reconstructs the input data from the latent vector. The models are trained to maximize the ability to reconstruct the input data with minimal information loss in the latent vector encoding. The logic behind using these for anomaly detection is that the AE learns to reconstruct common events more accurately, as they are the supermajority of the training set, and produces a larger reconstruction error for rare events. When compared with techniques that directly compute the probability, these techniques are computationally efficient but lack the underlying rigorous justification.

This issue can be addressed by training a special type of AE called the denoising autoencoder (DAE) and using its reconstruction error as a metric for rarity. DAEs are designed to reconstruct the original data from a noisy version of the data; it can be shown that the reconstruction error for a DAE approximates the magnitude of the score function (the gradient of the logarithm, *∇* log(*p*)) of the probability density function (41) for the data distribution. For most density functions, the magnitude of the score function is small in regions where the probability mass is concentrated (high-density regions) and large in the low-density regions. The score function, therefore, is a good measure of the rarity of an event (see Figure 1 for example). Motivated by these arguments, in this study we employ a DAE for detecting rare events.

**Fig. 1.**
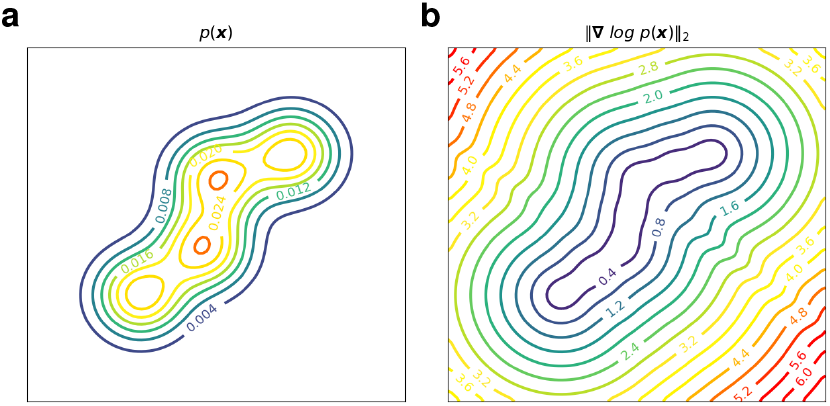
(a) Iso-contours of the probability density function (pdf) of a Gausian mixture model in two dimensions. (b) Iso-contours of the magnitude of the score function for the same pdf. Note that the score function is large in regions where the density is small.

Our approach begins by dividing a single four-channel immunofluorescence (IF) image of a slide into approximately 2.5 million tiles (see Figure 2). The size of the tile is selected so that each tile contains, on average, up to 4 events, where an event may be a cell, a vesicle or some other blood-based analyte. For applications considered in this study, this yields tiles with 32 × 32 pixels. Thereafter, uncorrelated Gaussian noise is added to each tile and pairs of clean and noisy tiles are used to train a DAE. When the training is complete, each tile is used as input to the DAE and the magnitude of the difference between the output of the DAE and the tile itself is evaluated for each IF channel. This scalar is multiplied with user-supplied channel weights, such that markers with important variance in the assay are emphasized, and the resulting products are summed to yield a single reconstruction error value for each tile. This error is used as a rarity metric to rank the tiles from most rare (largest reconstruction error) to least rare and a cohort 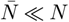 rare tiles is identified. In the final step, an algorithm to remove imaging artifacts from the rare tile cohort is applied and tiles with artifacts are replaced with tiles with slightly lower rarity metric. The approach is described in detail in the Methods section. We refer to this algorithm as the Rare Event Detection algorithm, or the RED algorithm in short.

**Fig. 2.**
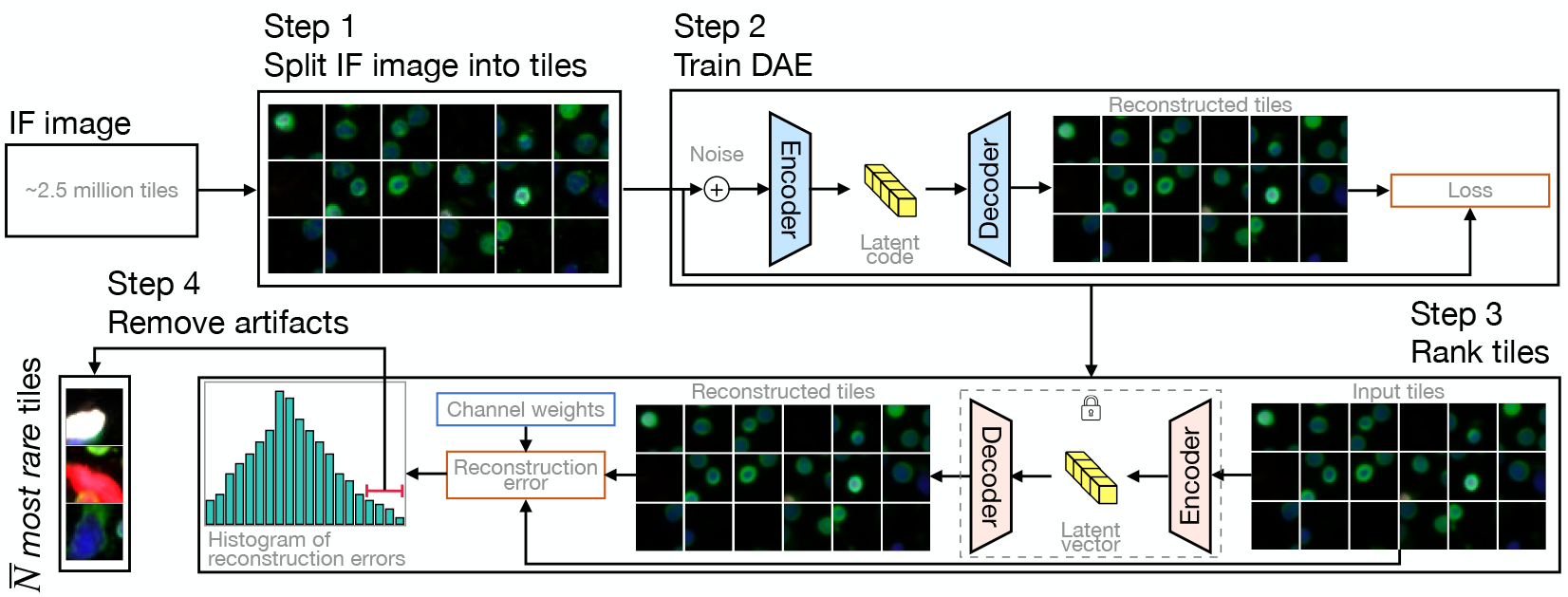
Schematic diagram of the rare event detection (RED) pipeline. In Step 1, on IF image is split into *≈* 2.5 million non-overlapping tiles. In Step 2, pairs of synthetically generated noisy tiles and their clean counterparts are used to train a denoising autoencoder (DAE). In Step 3, noisy tiles are used as input to the trained DAE and the difference between the de-noised and the original clean version of the tiles is used in combination with user-specified IF channel weights to evaluate the reconstruction error for each tile. Tiles with large values of the reconstruction error are identified and are deemed as being rare. In Step 4, an approach that assumes that true rare events are unlikely to be localized to a region within an IF image is used to eliminate artifacts from the rare tile cohort.

## 2. Results

In this section we describe the results obtained from applying the RED algorithm to two sets of IF images. The first set corresponds to blood from normal donors that is spiked with two different cell types, while the second set corresponds to blood from late stage breast cancer patients. Both sets comprise IF images with four channels representing DAPI (for DNA), a cocktail of cytokeratins (for epithelial cells) labeled with Alexa Fluor 555, vimentin (for mesenchymal cells) labeled with Alexa Fluor 488, and CD45/CD31 (for immune and endothelial cells, respectively) multiplexed in the same channel, labeled with Alexa Fluor 647. In order to keep the notation succinct, we refer to these channels as D, CK, V and CD, respectively. The collection and preparation of the samples, the construction of the assay, and the image acquisition are described in Section 4.1. The subsequent steps that begin with an IF image for a given subject and end with the rank ordering of each tile (defined as a 32 × 32 × 4 sub-region of an image) as per its rarity metric are described in Section 4.2.

In order to assess the utility of the RED algorithm, we adopt the following perspective. We note that a typical IF image contains around *N* ≈ 2.5 million tiles, and most of these contain immune cells that are not biologically interesting. Our hypothesis is that the RED algorithm is able to reduce this number down to a cohort that is about a thousand-fold smaller, 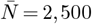, without eliminating a significant proportion of biologically interesting cells. We note that the utility of the much smaller rarity-ranked cohort is that it enables manual and automated downstream tasks, including single cell genomics and proteomics, that would not be feasible when working with the original cohort of 2.5 million tiles. Further, it is likely that there is utility in the ranking itself - that is, the fact a tile appears higher in the ranking is likely to be significant - though this remains to be verified in later studies.

For a given value of 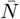, the rare tile cohort identified by RED represents tiles that have been classified as containing an interesting event. In order to quantify the performance of this classification, we compare this set with an independent set that is determined through an alternate, human-assisted pipeline described in our earlier work (7, 25, 42) and summarized in Section 4.2. We refer to this approach as the Outlier Clustering Unsupervised Learning Automated Report (OCULAR) pipeline. In this pipeline, several machine learning algorithms are first used to identify an average of approximately 3,000 (1,172 to 10,617, *M* = 3, 162, *SD* = 2, 676) potentially interesting events in an IF image. This is followed by a step where two human-trained analysts select the biologically interesting events from this reduced set. We treat the set identified by the OCULAR pipeline as the reference, and report our true positive rate (TPR) relative to this set. We also vary 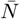 and construct the receiver operator characteristic (ROC) curve for our approach. We plot the ROC curve and report the area under the curve (AUROC), noting that only the initial part of the curve, where 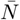 is small, is useful in an application of the RED algorithm.

For the late stage breast cancer patients, we also quantify the performance of the RED algorithm using a human-assisted pipeline. Within this pipeline, the 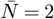, 500 rare tiles identified by the RED algorithm for every subject are examined by two human experts, who extract the biologically interesting events from this cohort. We do this to identify events that were detected by the RED algorithm but not the OCULAR pipeline. There are two important metrics to assess the performance of the RED algorithm: the percentage of events detected by the OCULAR pipeline that are also detected by the RED algorithm and the number of the additional events that are detected by the RED algorithm. We find that the RED algorithm finds 66 out of the 79 events detected by OCULAR; additionally it finds 91 events that are not detected by the OCULAR pipeline. This, along with the fact that it requires minimal manual optimization, clearly illustrates the utility of this approach.

### 2.1. Rare event detection in spiked cell samples

The ND samples with cell lines (SK-BR-3 and HPAEC cell lines) spiked in comprise nine IF slides. Of these nine slides, three are spiked with only SK-BR-3 cells, three are spiked with only HPAEC cells, and three are spiked with both. The SKBR-3 cells are a model system for rare epithelial cells or CTCs, while the HPAEC cells are a model system for rare endothelial cells. On average, each IF slide contains 342 (min. = 19, max. = 1030) spiked-in cells as identified by the OCULAR pipeline.

For each IF image we apply the RED algorithm, vary 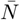 from zero to *N* and compute the ROC curves consisting of the FPR and TPR values for each 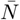. We do this for each spiked cell type separately and also for both cell types combined. This results in six ROC curves for SK-BR-3, six ROC curves for HPAECs, and nine ROC curves both cell types combined. For each set (SK-BR-3, HPAEC cells, and combined) we evaluate the lower quartile, median, and upper quartile ROC curve values. In Fig. 3 we plot the initial part of these curves (until FPR = 0.001). The solid curve represents the median, and the dashed lines represent the lower and upper quartiles. We observe that in every case the new algorithm yields a mean TPR close to unity (0.993, 0.965, and 0.985) for a very small FPR = 0.001. We do not plot the entire ROC curve since the values of the area under the ROC curve (AUROC), which is reported in Table 1, are very close to 1 and these curves do not reveal any information beyond this.

**Table 1.** Application of the rare event detection algorithm to the spiked cell dataset. Columns 2-5: statistics for the true positive rate for a cohort of 2,500 rare tiles identified by the algorithm. Columns 6-9: statistics for the AUROC obtained by varying 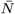 from 0 to *N*. Values are reported for CTCs (Row 1), endothelial cells (Row 2) and their combination (Row 3).

| Cell type | TPR ($\bar{N} = 2,500$) | | | | AUROC | | | |
| --- | --- | --- | --- | --- | --- | --- | --- | --- |
|  | Mean | St. dev. | Min. | Max. | Mean | St. dev. | Min. | Max. |
| SK-BR-3 | 0.993 | 0.014 | 0.962 | 1.00 | 1.00 | 0.0 | 1.00 | 1.00 |
| HPAEC | 0.965 | 0.031 | 0.927 | 1.00 | 0.999 | $2.00 \times 10^{-3}$ | 0.993 | 1.00 |
| All | 0.985 | 0.020 | 0.943 | 1.00 | 0.999 | $1.00 \times 10^{-3}$ | 0.997 | 1.00 |

**Fig. 3.**
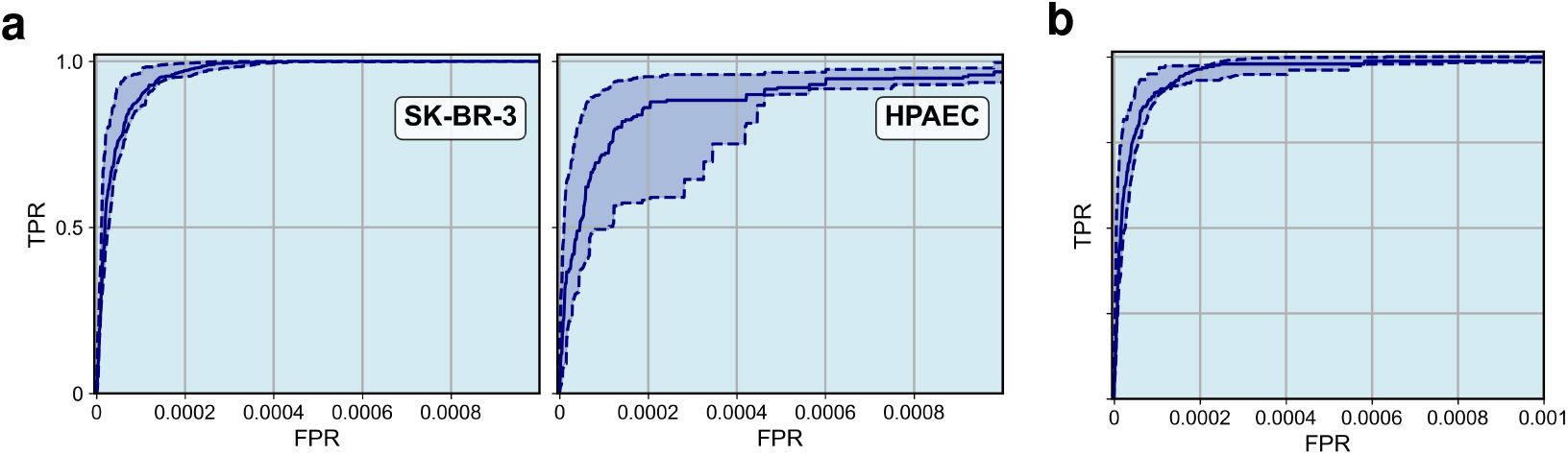
Initial part of the ROC (FPR range from 0 to 0.001) curve for the rare event detection algorithm applied to the spiked cell slides. Subfigure (a) shows the ROC curve for SK-BR-3 and HPAEC cell lines separately, while subfigure (b) shows the ROC curve for both cell lines. The solid curves represent the median ROC across all subjects, and the dashed curves represent the lower and upper quartiles.

In Table 1, we report the statistics for TPR across the nine subjects for 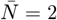, 500 noting that this value of 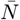 corresponds to a 1,000-fold reduction in data. For both cell types, the value of TPR with this 1,000-fold reduction in data is high (mean = 0.993 for SK-BR-3 and mean = 0.965 for HPAEC). Overall, with 1,000-fold data reduction using the RED algorithm we miss around 1.5% of biologically relevant events. In this table we also report the area under the ROC curve (AUROC) for the two cell types and all spiked cells taken together. The AUROC values obtained are very close to unity. In Fig. 4 we plot some of the tiles from the two spiked cell lines (SK-BR-3 and HPAEC) that were detected by the RED algorithm within a cohort of 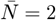, 500 tiles. We also plot some that were missed. We observe that the tiles that were detected tended to contain large, bright pieces of relevant cells, whereas those that were missed contained smaller pieces.

**Fig. 4.**
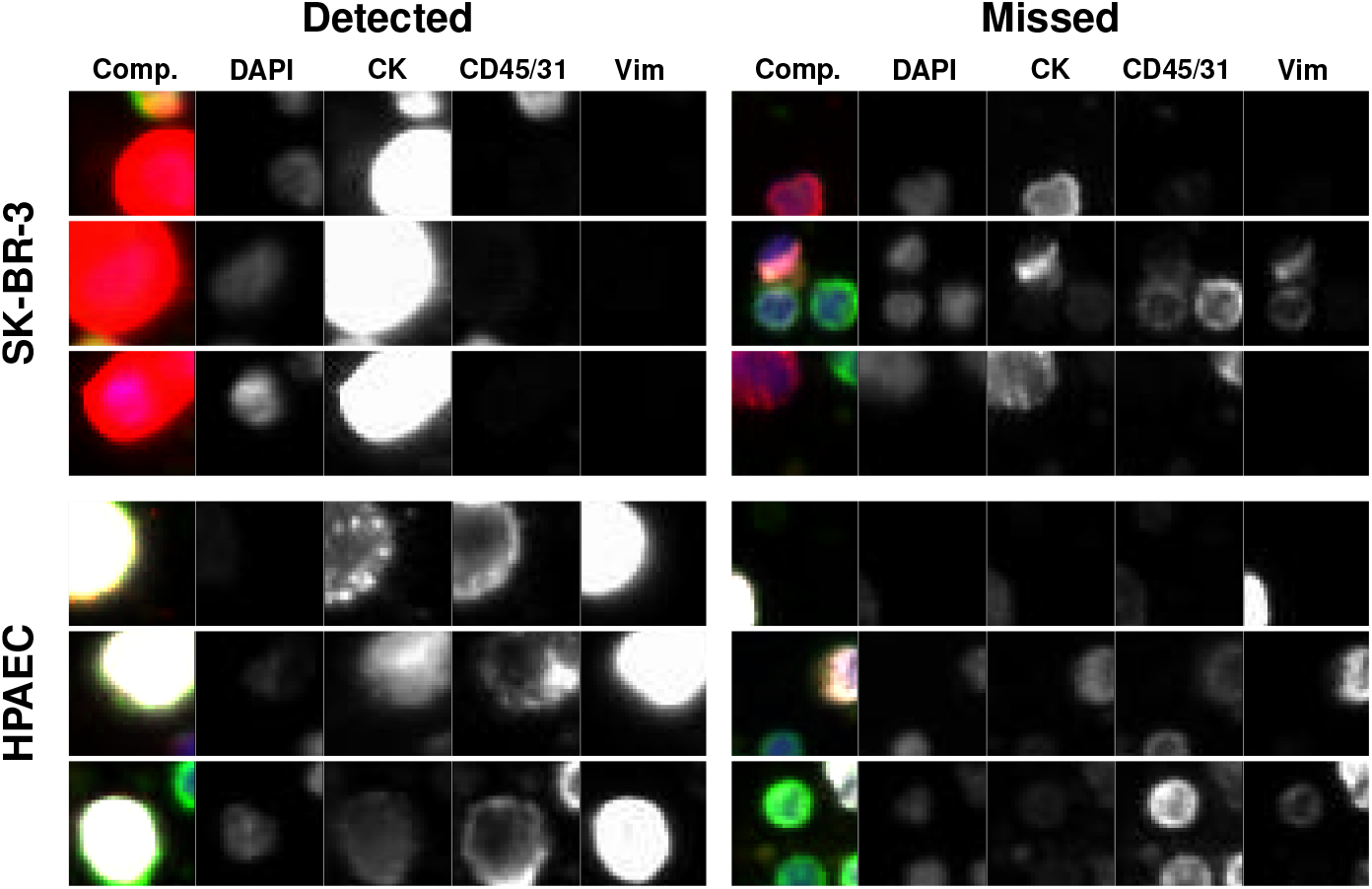
Representative gallery of rare events in samples from normal donors spiked with SK-BR-3 and HPAEC cell lines. For each rare event the composite image is shown followed by the biomarker fluorescent channels (specified by the headers). The top three rows show SK-BR-3 events and the bottom three rows show HPAEC events. The left column shows the events detected by RED and the right column rows shows the events not detected by RED.

### 2.2. Detection of rare cells in breast cancer patients

The late-stage breast cancer set comprises eleven IF labeled slides with each slide representing a sample from a unique late-stage breast cancer patient. On average each IF slide contains 8 (min. = 2, max. = 14) biologically relevant events as identified by the OCULAR pipeline. These biologically relevant events can be grouped into seven categories based on signal in the following channels: D-|CK, D|CK, D|CK|V, D|CK|V|CD, D|V, D|V|CD, and D|CK|CD, where D-denotes a DAPI negative signal, indicating acellularity.

We apply the RED algorithm to these images, and in Figure 5, plot the lower quartile, median and upper quartile ROC curves obtained by varying 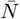 across all subjects and for the seven event categories, as well as all categories combined. The solid curve represents the median ROC curve while the dashed curves represent the upper and lower quartile variations about the median. In Figure 6, we focus on the earlier part of the ROC curves (FPR = 0.001). We observe that the performance of the RED algorithm for this set is not as good as for the spiked cell set (AUROC = 0.982 across all cell types). Further, there is significant variability in the performance across different event categories. From Figure 6 we observe that the algorithm performs well for some event categories (e.g., D|CK, D|CK|V, and D-|CK positive events) and is challenged in detecting others (e.g., D|V positive events). In Table 2, we have reported the TPR for the RED algorithm with a thousand fold reduction in data 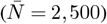. We observe that for a thousandfold data reduction, the median TPR across all event categories is 0.746, which is lower than the corresponding value for the spiked cell set. This can be attributed to the uncertainty in defining what constitutes a biologically relevant event in cases where these events occur naturally (as in the late-stage breast cancer set) and are not introduced artificially (as in the spiked cell set). This makes the detection of these events difficult for the RED algorithm as well as OCULAR pipeline, which is the approach used as the reference.

**Table 2.**
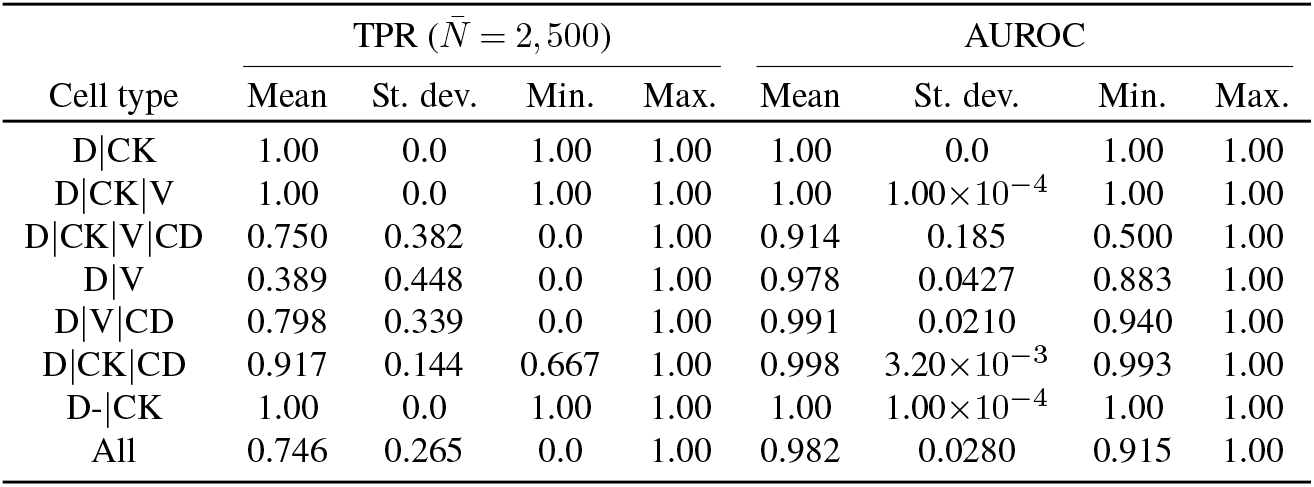
Application of the rare event detection algorithm to images from late-stage breast cancer subjects. Columns 2-5: statistics for the true positive rate for a cohort of 2,500 rare tiles identified by the algorithm. Columns 6-9: statistics for the AUROC obtained by varying 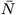 from 0 to *N*. Values are reported for different event types (Rows 1-7) and all types together (Row 8).

**Fig. 5.**
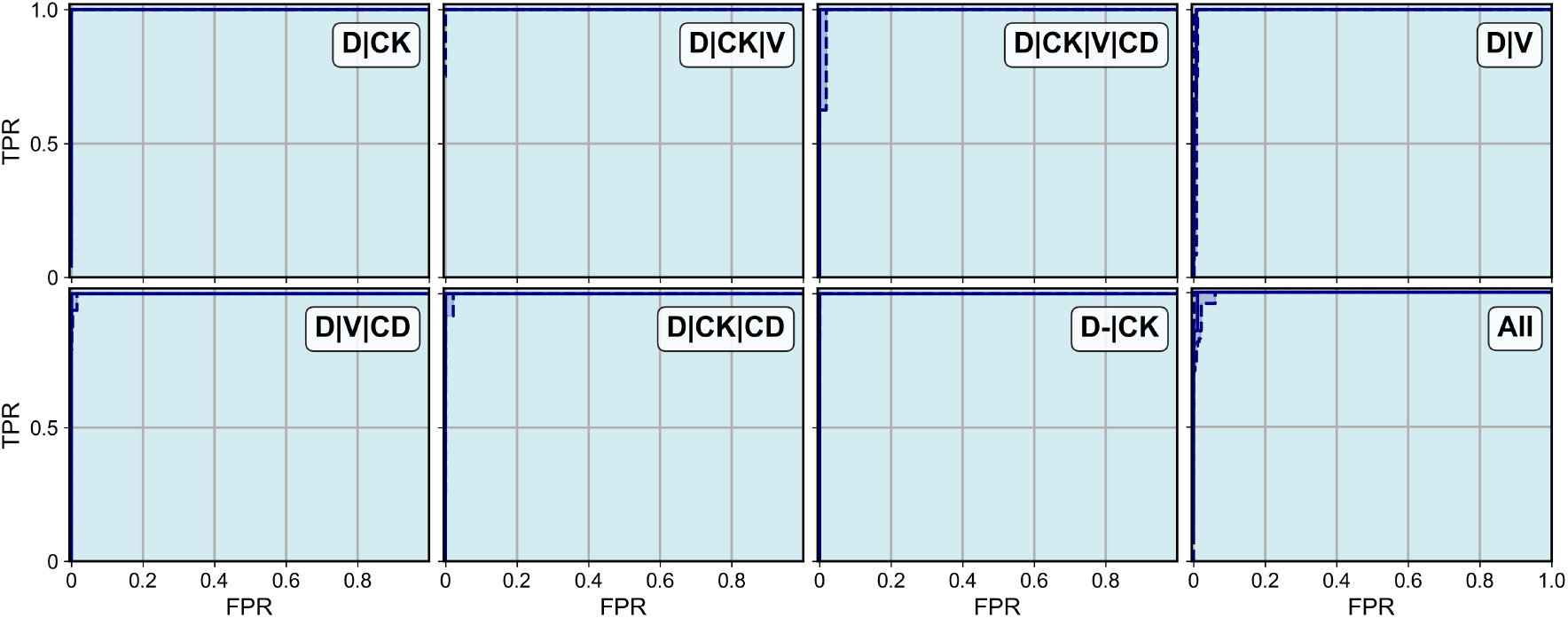
ROC curves for the rare event detection algorithm applied to late stage breast cancer subjects. Separate ROC curves are shown for each event type as well as all event types combined (bottom right). The solid curves represent the median ROC across all subjects, and the dashed curves represent the lower and upper quartiles.

**Fig. 6.**
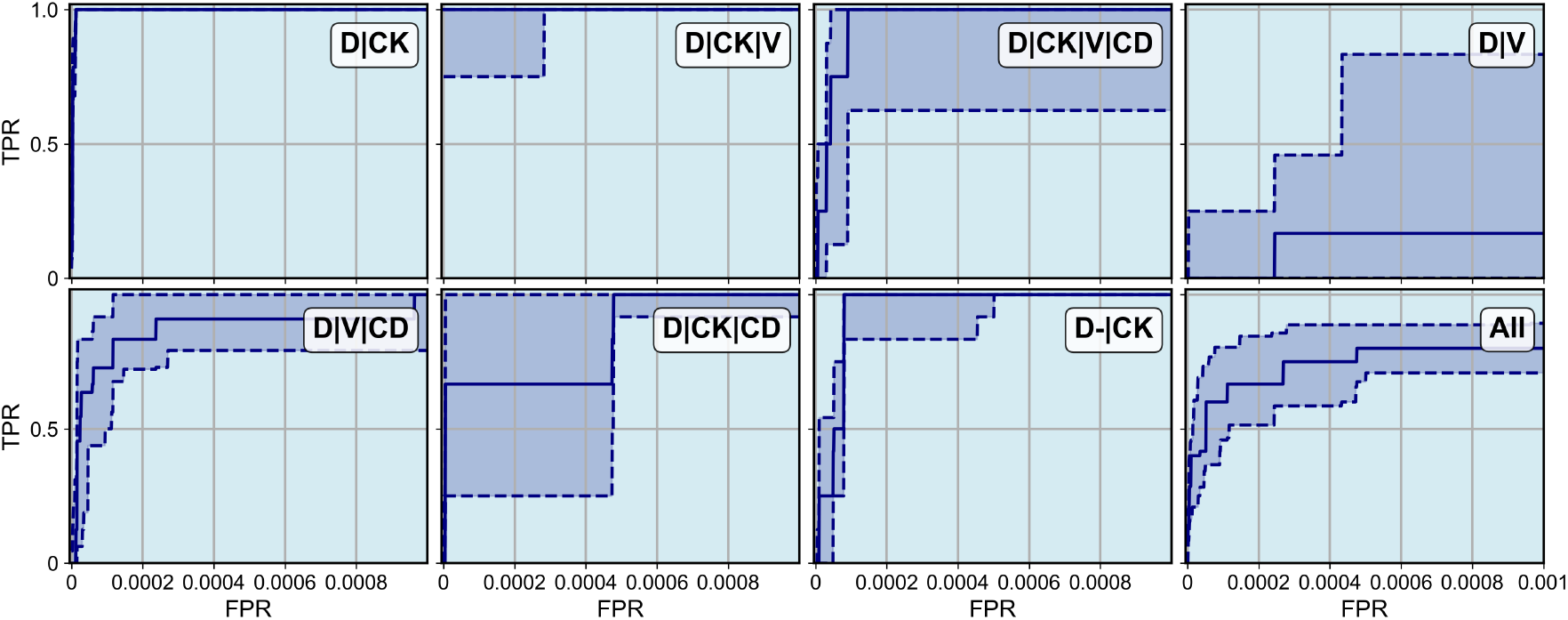
Initial part of the ROC curves for the rare event detection algorithm applied to late stage breast cancer subjects. Separate ROC curves are shown for each event type as well as the composite ROC curve for all event types combined (bottom right). The solid curves represent the median ROC across all subjects, and the dashed curves represent the lower and upper quartiles.

Fig. 7 shows a sample of the tiles from the late-stage breast cancer slides that were detected by the RED algorithm within a cohort of 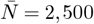 tiles and some tiles that were not detected within that cohort. In three out of the seven categories we report no missed tiles.

**Fig. 7.**
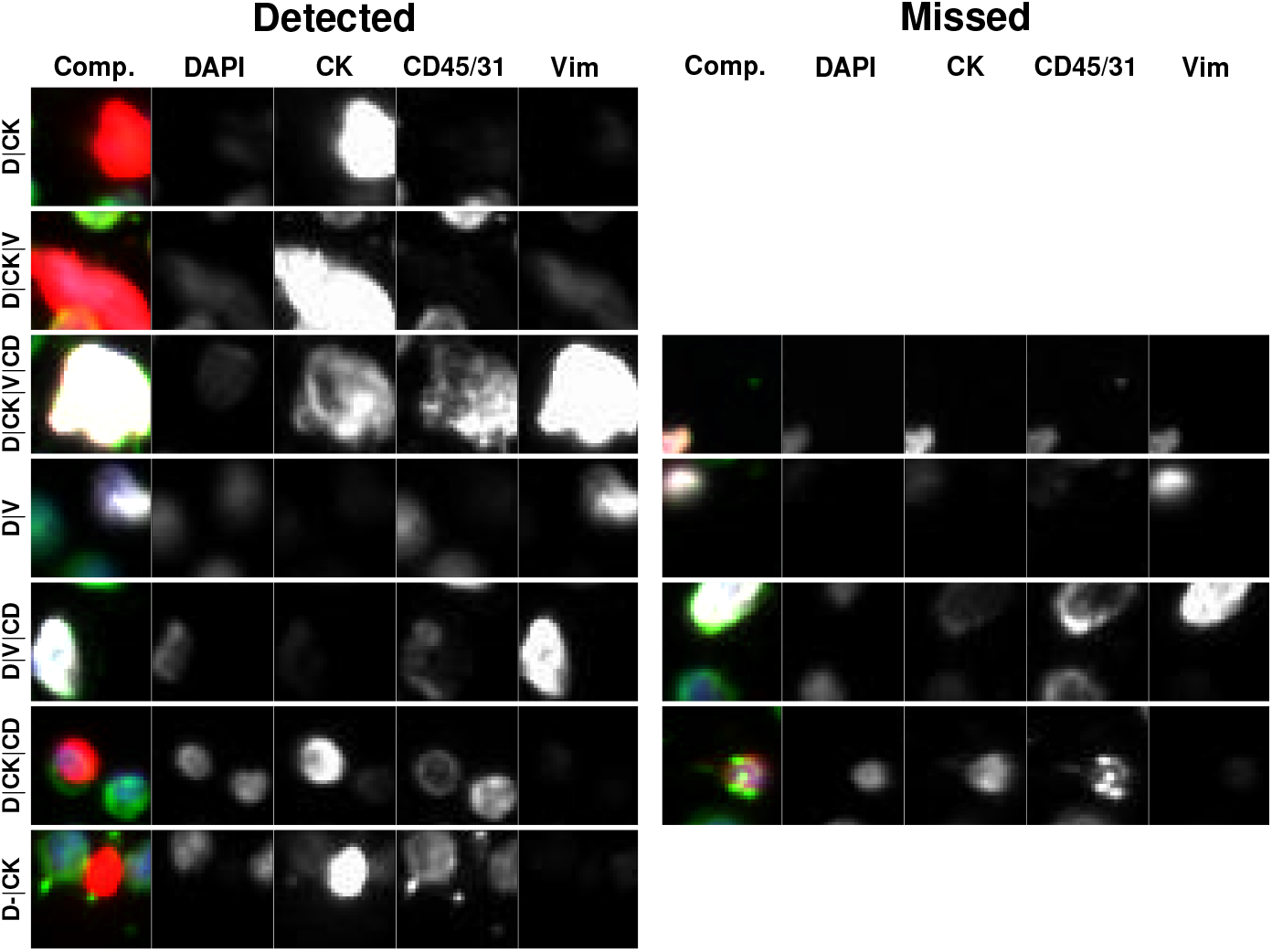
Representative gallery of rare events in samples collected from patients diagnosed with late-stage breast cancer. For each rare event the composite image is shown followed by the biomarker fluorescent channels (specified by the headers). The left column shows rare events detected by RED and the right column shows rare events not detected by RED. No event is shown for the cell types for which no event was missed.

A manual examination of the set of 2,500 events identified by the RED algorithm revealed that this set included several events that were biologically relevant but were not identified by the OCULAR pipeline. In hindsight, we should have anticipated this since the OCULAR pipeline also has its own false negative errors. This realization led us to consider the approach described below for quantifying the performance of the RED algorithm.

As described in Section 4.2, the OCULAR pipeline consists of two distinct stages. In the first stage, all events in a given IF slide are segmented and a short-list comprising approximately 3,000 interesting events is identified by the OCULAR algorithm. Events in this short-list are then examined by multiple human experts and those deemed to be biologically interesting by both experts are included in the final list of biologically relevant events. Analogous to this, we develop and implement the RED pipeline where the 2,500 events per IF slide identified by the RED algorithm were examined by two human experts, and those deemed to be biologically interesting by both experts are included in the final list of biologically relevant events binned into one of the seven event categories defined above.

Once the OCULAR and RED pipelines have identified the set of biologically relevant events, we computed the number of events detected by both pipelines and each pipeline alone. These numbers are reported in Figure 8. We observe that the RED pipeline identifies around twice as many events when compared with the OCULAR pipeline (157 versus 79). Another way to measure the efficacy of the two pipelines is to consider the number of events identified by only one pipeline. In this respect the RED pipeline identifies seven times as many events as the OCULAR pipeline (91 versus 13). We note that the performance of the RED pipeline is dependent on the event category. In particular, for the D-| CK category the RED pipeline identifies around 8 times as many events as the OCULAR pipeline(73 vs. 9), while for D|V events the OCULAR pipeline performs slightly better (11 vs. 12).

**Fig. 8.**
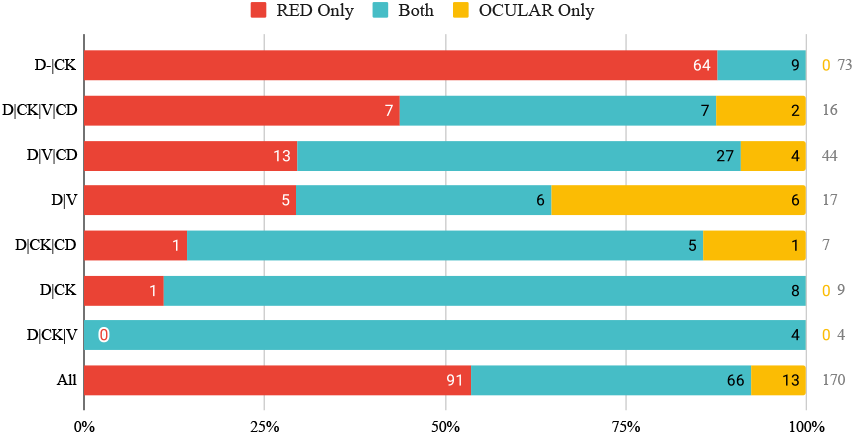
Enumeration of biologically relevant events identified by the RED pipeline alone (in red), the OCULAR pipeline alone (in yellow) and both pipelines (in blue). Rows 1-7 depict results for 7 different event types, while row 8 depicts composite results for all event types.

## 3. Discussion

The RED algorithm represents a paradigm shift in detecting biologically relevant events in LBx. Most current methods seek specific analytes in LBx assays through physical enrichment. This can be challenging when there is not a single analyte of interest but rather a heterogeneous population. Further, in exploratory studies where the analyte of interest is not known, it is impossible to use these types of methods. In contrast, the RED algorithm works on the simple premise that biologically relevant information is rare relative to the common immune population. This obviates the need to specify the characteristics of what constitutes a biologically relevant event and makes the detection task simpler and easier to automate.

When compared with the baseline approach (OCULAR algorithm), the RED algorithm comprises fewer steps that are easier to automate and require minimal expert guidance. In particular, in the RED algorithm, the steps required to get to the cohort of 2,500 rare events are: training the DAE, using the DAE to rank tiles, and removing artifacts through an automated approach. The expert input required for these steps is limited to specifying the channel weights (four scalar values), and the threshold (a single scalar value) used in removing tiles that contain artifacts. In contrast, the steps in OCULAR include threshold segmentation of around 2.5 million events, evaluation of 761 parameters for each event, reduction of these to 350 PCA components, and cascading clustering stages which result in a wide range of events retained for final analysis. These steps require the specification of: (a) hyperparameters for segmentation, (b) each of the 761 features to be computed for each event, (c) number of PCA components to be retained, (d) majority cluster elimination from the cascading cluster stages to help with negative depletion of majority class and (e) distances that constitute an adequately rare event compared to median references. Overall, this requires significantly more information to be specified by a computational expert, which makes this approach harder to automate. Further, the RED algorithm retains only 2,500 rare events per IF slide, whereas the OCULAR algorithm retains roughly 3,000 events per IF slide, both of which require some level of human data curation. As such, the RED algorithm leads to more significant data reduction, which makes downstream analysis easier and more efficient.

In the spiked cell cohort considered in this study, most of the epithelial and endothelial cells were captured in the set of 2,500 rare tiles identified by the RED algorithm. On average it missed 0.7% of the epithelial cells and 3.5% of the endothelial cells. This served to validate the performance of RED in a case where the biologically relevant events were well known and could be easily characterized.

The late stage breast cancer cohort comprised naturally occurring biologically relevant events that were not contrived. In this case, the rare tiles identified by the RED algorithm were examined by two experts in order to select biologically interesting events. The performance of this pipeline, which was dubbed the RED pipeline, was compared with that of a similar analysis which used OCULAR to identify the rare events. It was found that the RED pipeline yielded twice as many biologically relevant events, which points to its utility in real-world applications. It was also found that the RED algorithm was able to detect most of the events detected by the OCULAR pipeline (84%, Figure 8).

Additionally, it is clear that the performance of the RED pipeline relative to the OCULAR pipeline varies depending on the event category in question. For the samples considered in this study, RED performed significantly better than OCULAR for D|CK events, and slightly worse than for D|V events. This may be attributed to the fact that CK positive events, whether they are biologically relevant or not, are rare in the peripheral blood context, and can therefore be easily identified by a rarity detection algorithm like RED. In contrast to this, V-positive events are common, with a population of leukocytes expressing vimentin, and a very small fraction of these cells (V-positive with variable expression in the other channels) are biologically relevant. In this case a rarity detection algorithm has to work “harder” relying on factors like cell morphology and relative intensity across multiple channels in order to detect biologically relevant rare events. Overall, the analysis shows that the RED pipeline performs better than the baseline method OCULAR for every event category except D|V where its performance is marginally worse (11 versus 12 events identified).

The RED methodology offers improved sensitivity over the baseline OCULAR method. RED identifies a greater number of rare events, which is critical for enhancing detection capabilities in a rarity-focused framework. This methodology is particularly advantageous because it is largely automated, reducing human involvement and thereby minimizing potential sources of error and the time required for analysis. Despite these improvements, molecular characterization of the detected events is essential to elucidate their biological and clinical relevance. For instance, D|CK cells are consistent with canonical epithelial CTCs, and D|V|CD cells are morphologically and phenotypically consistent with circulating endothelial cells. Additionally, the D-|CK events identified by RED are hypothesized to be oncosomes or large extracellular vesicles potentially associated with the disease state (43). Further validation studies will confirm these biological phenotypes and provide deeper insights into their role in cancer biology and the potential clinical implications.

When compared to the CellSearch platform, which is a widely used enrichment-based approach clinically utilized in breast cancer patient care, RED demonstrates distinct advantages. CellSearch is tailored to detect known cell types, specifically circulating tumor cells that are EpCAM+, CK+, and CD45-, and relies on a predefined set of markers. While effective for certain applications, this targeted approach introduces bias and limits the detection of rare and unconventional events, such as oncosomes or tumor microenvironment components like endothelial cells or fibroblasts. In contrast, RED’s unbiased framework allows for the identification of a broader range of rare events, enabling novel discoveries and expanding the potential applications of liquid biopsy. These attributes position RED as a transformative tool in rare event detection, with the capacity to uncover previously undetected facets of disease biology.

Single-channel biophysical enrichment approaches, while streamlined, often result in the loss of multidimensional enrichment capabilities, which are crucial for capturing the complex heterogeneity of rare events. This limitation underscores the importance of a methodology like RED, which preserves sensitivity to rare populations without compromising the breadth of detection. Given the rare event framework (employed by the HDSCA platform) has demonstrated more sensitivity than CellSearch in detecting cellular heterogeneity and plasticity (44–46), and RED shows improved sensitivity and detection capabilities beyond the OCULAR workflow used by HDSCA, then it represents a clear step forward in the evolution of rare event detection technologies. Moreover, RED offers a distinct advantage from a development perspective. Its algorithmic design simplifies the process of enriching rare event populations, making it “lightweight” and user-friendly for developers. This reduced dependency on deep biological understanding allows researchers to focus on refining detection and analysis pipelines rather than grappling with complex enrichment processes. As a result, RED provides a robust, scalable framework that maximizes detection sensitivity and operational efficiency, facilitating both innovative discoveries and ease of adoption in research and clinical settings.

## 4. Methods

### 4.1. Blood collection, sample preparation and imaging

Peripheral blood (PB) samples were collected in cellfree DNA blood collection tubes (Streck, La Vista, NE USA) and processed as previously described (44, 45, 47). Briefly, after complete blood cell count (Medonic M-series Hematology Analyzer, Clinical Diagnostic Solutions Inc., Fort Lauderdale FL USA) the red blood cells were lysed with ammonium chloride and all nucleated cells were plated as a monolayer on custom cell adhesion glass slides (Marienfeld, Lauda, Germany) at approximately 3 million cells per slide, followed by blocking with 7% bovine serum albumin (BSA) before drying and cryopreservation at -80 ° C.

Samples were stained automatically (IntelliPATH FLX autostainer, Biocare Medical LLC) with the Landscape immunofluorescence (IF) assay as previously published(5, 9, 25, 42, 48–50). Briefly, slides were thawed and fixed with 2% paraformaldehyde prior to 1) incubation with antihuman CD31 Alexa Fluor 647 direct conjugate (mouse IgG1 monoclonal antibody; 2.5 *µ*g/mL; clone: WM59; Cat# MCA1738A647; BioRad; RRID:AB 322463) and anti-mouse Fab fragments (IgG goat monoclonal; 100 *µ*g/mL; Cat# 115–007–003; Jackson ImmunoResearch), 2) permeabilization with cold methanol, 3) incubation with a mixture of anti-human pan cytokeratin (CK) (CKs 1,4,5,6,8,10,13,18,19 mouse IgG1/IgG2a monoclonal antibody cocktail; 210 *µ*g/mL; Cat# C2562; clone: C-11, PCK-26, CY-90, KS-1A3, M20, A53-B/A2; Sigma; RRID:AB 476839), anti-human CK 19 (mouse IgG1 monoclonal antibody; 0.2 *µ*g/mL; Cat# GA61561–2; clone: RCK108; Dako), anti-human CD45 Alexa Fluor 647 direct conjugate (mouse IgG2a monoclonal antibody; 1.2 *µ*g/mL; Cat# MCA87A647; clone: F10–89–4; AbD Serotec; RRID:AB 324730), and anti-human vimentin (VIM) Alexa Fluor 488 direct conjugate (rabbit IgG monoclonal antibody; 3.5 *µ*g/mL; Cat# 9854 BC; clone: D21H; Cell Signaling Technology; RRID:AB 10829352), and 4) incubated with anti-mouse IgG1 Alexa Fluor 555 (goat IgG polyclonal antibody; 2 *µ*g/mL; Cat# A21127; Invitrogen; RRID:AB 141596) and 4’,6-diamidino-2-phenylindole (DAPI; dilution: 1: 50,000; Cat# D1306; Thermo Fisher Scientific; RRID:AB 2629482). Slides were mounted with a glycerol-based media, coverslipped, and sealed.

Automated scanning was done at 100X magnification using a custom high-throughput fluorescence scanning microscope across 2304 frames per slide in each channel (DAPI, Alexa Fluor 488, Alexa Fluor 555, Alexa Fluor 647). The exposure time and gain per channel were automatically set to ensure consistent background intensity across all slides for normalization purposes.

Normal donor (ND) samples were procured from the Scripps Normal Blood Donor Service and processed according to the above. Cell line cells with known expression profiles were spiked into the sample at various concentrations after red blood cell lysis (SK-BR-3 ATCC HTB-30 and HPAEC ATCC PCS-100-022). Standard protocols were followed for contrived sample analysis.

A total of 11 samples collected from patients with metastatic breast cancer were included in this study. Patient recruitment took place according to an institutional review board-approved protocol approved by the University of Southern California (FWA 00007099, USC UPIRB #UP-14-00523) and all study participants provided written informed consent (44, 51).

### 4.2. Rare event detection (RED) algorithm

In order to detect rare events within the IF assay, we employ a deep learning method for anomaly detection. In the first step of our approach we split the IF image for a subject into a set of non-overlapping sub-images that we refer to as tiles (see Figure 2). The size of a tile is selected so that each tile includes 1-4 cells on average. In our case, this corresponds to a size 32 by 32 pixels, or 18.9 by 18.9 *µ*m, which yields approximately 2.5 million tiles per IF image.

The collection of tiles generated is used to train a denoising autoencoder (DAE). The architecture, training-related hyper-parameters, and the loss function used for training the DAE are reported in Appendix A. During training, the input to the DAE is a noisy version of each tile and its output is the corresponding de-noised version. The noisy version of a tile is generated by artificially adding uncorrelated homoscedastic Gaussian noise (with variance = 0.3^2^) to every pixel of the tile. The DAE learns how to reconstruct tiles that contain common events well, but not tiles that contain rare events. Consequently, when tiles with common events are passed through the fully trained DAE it produces images that are close to the original tile, whereas for tiles with rare events this is not the case. The magnitude of the difference between the reconstructed tile and the original tile is computed on a per-channel basis. This magnitude is then multiplied with a channel-dependent weight and all the weighted values are added to arrive at a single real-valued reconstruction error, which is used as the rarity metric. The values of weights used in this study are 1/3 for the DAPI, CK and V channels, and 0 for the CD channel. Note that the DAE does not use any labeled data during training the DAE or when computing the rarity metric for each tile. Thus, our approach is unsupervised and works without any apriori information regarding biologically relevant events, such as location in the IF assays or phenotype, specific to the disease.

Tiles with large values of the reconstruction error are deemed as rare, where those with small values are deemed as being common. There is a theoretical justification of this observation. It can be shown that for a given input sample, the reconstruction error is an approximation to the magnitude of the score function of the underlying probability density for that sample (41). Further, since for most probability densities the magnitude of the score function is much larger in regions where the probability mass is small, the reconstruction error may be used as a metric for rarity.

During the application of the proposed approach we observed that some tiles that contain imaging artifacts were selected in the rare tile cohort. This is not surprising given the understanding that certain types of imaging artifacts can also be rare. In the examples considered in this manuscript the artifacts include speck-like regions with a strong signal in CK channel, and blurs and streaks across all channels. Both these artifacts tended to occur in clusters at a specific location of the image, and this characteristic was used to remove the tiles with these artifacts. For the specklike artifacts, the number of artifacts occurring within a sub-domain of an image was counted, and if this number exceeded a specified threshold, all tiles in the rare event cohort from that subdomain were removed. This subdomain was set to 1362 by 1004 pixels, the original size of the images taken by the scanning microscope, and the threshold used was 500 specks per subdomain. To eliminate other regionally concentrated artifacts present in the rare tile cohort, the number of tiles from the top 10,000 rare tiles per subdomain was calculated. If this number exceeded 25, all tiles from that subdomain were removed. This approach is based on the observation that artifacts tend to be regionally concentrated, whereas biologically significant events are dispersed throughout the IF image. Hence, removing a few subdomains (typically less than 2% of the image) has negligible effect on the biological signal while effectively removing artifacts from the top of the ranking. Note that the subdomains used for artifact removal are predefined and are non-overlapping.

### 4.3. OCULAR rare event algorithm

In this study OCULAR was used as a reference to quantify the performance of the RED algorithm. OCULAR is a custom algorithm for rare event detection used in the high-definition single cell assay (HDSCA) workflow that uses image processing for feature extraction, dimensionality reduction, and unsupervised clustering (7, 25, 42). Namely, the “EBImage” package (EBImage 4.12.2) is used to segment the fluorescent images for every event across the slide, separating cells (expressing DAPI) from acellular components (not expressing DAPI). This is followed by feature extraction for each cell, generating 761 quantitative parameters across the 4 IF channels and paired combinations of each. A principal component analysis (PCA) transform is calculated and each cell’s morphometric data is projected onto the top 350 principal components. This reduction was shown to retain 99.95% of the original variance. Next, event-to-event distances for all cells in a given frame are calculated and ≈ 30 hierarchical clusters are generated. Thereafter, a cell is defined as rare if it belongs to the smallest clusters until the number of cells added by including a cluster exceeds 1.5% of all events on a frame or if it is far away from the median event in a frame. After this frame-based identification, frames are clustered into 10 bins based on their aggregated feature values. Events in each group are first compared internally, where rare events are filtered based on distance to common event clusters, and then further filtered with the same method when aggregated with the whole slide. Around 3,000 cellular events are initially identified as rare and potentially interesting.

In the OCULAR pipeline, the events identified by the OCULAR algorithm described above are examined by two experts and those deemed as being biologically relevant by both experts are retained.

To compare the algorithms, we sought to identify biologically interesting events found through each method. First, two human experts evaluated composite RGB images and single-channel grayscale images for the OCULAR results, determining whether the events were biologically relevant. Events that both experts agreed were relevant were retained for a total of 113 OCULAR events. Next, the 2,500 rarest tiles as identified by RED were examined by one human expert as both composite RGB images and single-channel grayscale images to determine an initial subset of 609 potentially interesting tiles. A further 29 tiles that corresponded to OCULAR events but were missed in this evaluation were added to the potentially interesting tiles. Both experts independently evaluated each tile for biological relevance and tiles rated as irrelevant by either or both experts were removed. Tiles that captured components of the same event were also deduplicated, leaving 166 events. Both experts then categorized all events, where disagreements were resolved by deference to one expert or selection of the majority class in cases of events found in both pipelines. Finally, events categorized as D and D|CD, which are often biologically semi-interesting, as well as one event categorized as D-|CK|V|CD, were removed from both pipelines, leaving 157 RED tiles and 79 OCULAR events. Figure 8 illustrates the events identified by both algorithms, as well as the overlap in identified events, separated by channel classification.

We evaluated the reliability of expert classifications using Cohen’s kappa, a measure for interrater reliability that accounts for chance agreement. This metric ranges from 0 to 1, with 0 indicating no agreement and 1 indicating perfect agreement. Based on all classified events, including D and D|CD events, we found *κ* = 0.775 ± 0.058 (95% CI), or moderate agreement (52). This level of agreement illustrates the level of difficulty even for expert human curators to identify cell phenotypes and events of interest.

## Declarations

This work was supported in part by the Ming Hsieh Institute for Research on Engineering-Medicine for Cancer (A.O., P.K., J.M.); National Cancer Institute, U01CA285013 (P.K., J.H., J.M.); Breast Cancer Research Foundation, BCRF-23-089 (P.K., J.H.); the Miriam and Sheldon G. Adelson Medical Research Foundation (P.K.); and the National Cancer Institute’s Norris Comprehensive Cancer Center (CORE) Support 5P30CA014089-40 (P.K., J.H., J.M.). This work also received support from the Vassiliadis Research Fund. The content is solely the responsibility of the authors and does not necessarily represent the official views of the National Institutes of Health.

## Data availability

All data discussed in this manuscript would be available upon request.

## Code availability

The code to train the DAE and rank tiles according to the rarity metric are available at https://github.com/jmurgoitioesandi/Unsupervised_RareCellDetection/tree/main/DAE_RCD_TF2. Standard Python and matplotlib methods were used to generate the visualizations.

## A. Denoising autoencoder training details

The autoencoder model consists of an encoder and a decoder, with each composed of convolutional, dense, pooling, and upsampling layers. The details of the encoder and decoder architectures are described in Table 3. Note, the dimensionality of the autoencoder’s latent vector was chosen to be 100. The layers in the architecture shown in Table 3 are described as follows. *Linear(in, out)* represents a fully connected layer with *in* input dimensions and *out* output dimensions. *Conv2D(in, out)* are 2D convolutional layers with a kernel size of 3, where *in* is the number of input filters and *out* is the number of output filters. *AveragePool2D(pool_size, stride)* is a downsampling layer with the specified pooling size and stride. *Dense-block(k, n)* refers to the block architecture in (53), with *k* input filters and *n* layers. *Upsample(size, size)* represents an upsampling layer with a scaling factor of *size. BN* denotes batch normalization.

**Table 3.** Autoencoder architecture: encoder and decoder layers.

| Encoder | Decoder |
| --- | --- |
| Conv2D(4, 32) - ReLU | Linear(100, 300) - ReLU - BN |
| Dense-block(32, 3) | Linear(300, 2048) - ReLU |
| Conv2D(32, 64) - AvgPool2D(2, 2) - ReLU | Reshape(2, 2, 512) |
| Dense-block(64, 3) | Conv2D(512, 256) - ReLU - Upsample(2, 2) |
| Conv2D(64, 128) - AvgPool2D(2, 2) - ReLU | Dense-block(256, 3) |
| Dense-block(128, 3) | Conv2D(256, 128) - ReLU - Upsample(2, 2) |
| Conv2D(128, 256) - AvgPool2D(2, 2) - ReLU | Dense-block(128, 3) |
| Dense-block(256, 3) | Conv2D(128, 64) - ReLU - Upsample(2, 2) |
| Conv2D(256, 512) - AvgPool2D(2, 2) - ReLU | Dense-block(64, 3) |
| Flatten | Conv2D(64, 32) - ReLU - Upsample(2, 2) |
| Linear(2048, 300) - ReLU - BN | Dense-block(32, 3) |
| Linear(300, 100) | Conv2D(32, 4) - Sigmoid |

The denoising autoencoder was trained using the mean squared error (MSE) loss function in Eq. S (1), as described in (54)

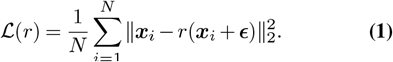

In this equation, 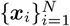 represents the set of input tiles, and *r*(***x***_*i*_ + ***ϵ***) denotes the autoencoder’s reconstruction of each input tile after adding noise. The noise, ***ϵ***, is sampled independently for each tile from a Gaussian distribution 𝒩 (**0**, 0.3^2^***I***), where ***I*** is the identity matrix. For each slide, an autoencoder is trained for 50 epochs using the Adam optimizer with learning rate equal to 10^*−*5^ and batch size equal to 500.

The DAE was trained on a NVIDIA V100 GPU, with each slide taking approximately 30 minutes to train.

